# Predicting gene-specific regulation with transcriptomic and epigenetic single-cell data

**DOI:** 10.1101/2025.11.16.688671

**Authors:** Laura Rumpf, Fatemeh Behjati Ardakani, Dennis Hecker, Marcel H Schulz

## Abstract

**Motivation:** Analysis of single cell ATAC-seq and RNA-seq data has allowed to gain unprecedented insights into gene regulation by allowing to define cell type specific regulatory regions and their effects on gene expression. While powerful, such analysis is challenging due to the inherent sparsity of single cell data.

**Results:** We present the MetaFR approach to learn gene-specific models that link open-chromatin variation from scATAC-seq data to gene expression from scRNA-seq. Using efficient regression trees, we illustrate that accurate expression prediction models can be learned on the single-cell or meta-cell level. Validation was done using fine-mapped eQTLs. Meta-cell models were found to outperform single-cell models for most genes. Comparison to the SOTA method SCARlink revealed advantages of MetaFR in terms of runtime and prediction performance. MetaFR thus allows time-efficient analysis and obtains reliable models of gene expression prediction, which can be used to study gene regulation in any organism for which scRNA-seq and scATAC-seq data is available.

**Availability and implementation:** MetaFR is freely available under https://github.com/SchulzLab/MetaFR.

## Introduction

A prevailing challenge in the field of epigenetics stems from the limited understanding of transcriptional regulation diversity. Cis-regulatory elements such as enhancers or silencers play a central role in gene regulatory programs that shape cellular development and identity ([17, 25, 36], [37]). They have been linked to several diseases and have been shown to be promising candidates for molecular targets in therapeutic interventions ([3, 29, 37]).

Genome-wide measurements of transcriptional and epigenetic activity have led to the development of numerous approaches for linking enhancers to genes and predicting gene expression. The integrative analysis of both modalities measured in individual cells enables cell type-specific insights into gene regulation mechanisms ([31]). Epigenome-expression modeling approaches can be divided into two categories: gene-agnostic and gene-specific ([27]). Gene-agnostic approaches model all genes jointly, regarding them as a uniform set of training instances. For instance, sequence-based approaches utilize the DNA sequence from a single biological sample as input to predict epigenetic signals, gene expression levels, or both, genome-wide ([1, 15, 16, 38]). By doing so, these approaches bypass the need for large numbers of biological samples, as they exploit the complete genetic information of the genome.

In contrast, gene-specific approaches leverage the variability of the epigenetic signal across samples usually within a window surrounding a gene to predict gene expression. In this case, one model per gene is learned to associate the epigenome with the gene expression ([8, 11, 13, 20, 21, 28, 30]). While gene-specific approaches are computationally intensive due to the large number of models that need to be trained, they are particularly well suited to harness the statistical power provided by the large number of samples in single-cell data. Accordingly, several gene-specific methods have been developed tailored for single-cell data.

In general, these approaches can be divided into univariate and multivariate ([8]). The univariate or correlation-based approaches model pairwise enhancer-gene associations ([11, 20, 26, 33]), while the multi-variate methods regress over all regions within a window around a gene ([8, 21]). The advantage of multivariate over correlation-based methods lies in their ability to capture co-regulatory effects, i.e., they can take into account the combined influence of several regulatory elements on gene expression rather than considering each element in isolation.

The state-of-the-art multivariate method SCARlink ([21]) utilizes multiome RNA-seq and ATAC-seq data to predict gene expression with regularized Poisson regression. One limitation of this method is its restriction to positive coefficients for easier model learning and interpretation. Consequently, SCARlink is not able to capture suppressive interactions, which are essential for transcriptional regulation ([9, 23]). [8] benchmarked SCARlink against other multivariate approaches for identifying enhancer-gene associations, showing that although SCARlink performed best, no method fully captures how chromatin accessibility regulates gene expression.

A particular challenge is the inherent sparsity of single-cell data. While a standard scRNA-seq matrix contains 55 − 95% zeros, scATAC-seq data is even more sparse with 90 − 99% zeros ([7]). As a result, extracting biologically meaningful insights remains challenging. A common strategy to address this problem is to aggregate the cells into meta-cells according to the similarity of their gene activity profiles ([4–6, 24]). [24] evaluated the effect of meta-cell aggregation using a correlation-based approach. However, for multivariate models, the impact of meta-cell aggregation on gene expression prediction accuracy remains unexplored.

We developed a fully automated Nextflow pipeline, **MetaFR** (Meta-Cell Forest Regression), that leverages single-cell RNA-seq and ATAC-seq data to build gene-specific regression trees for gene expression prediction. These trees avoid overfitting and are fast and we show that they outperform the state-of-the-art method SCARlink ([21]) in both accuracy and runtime. Further, we assessed the impact of meta-cell creation on model performance. Applying a signal aggregation strategy significantly improved the prediction accuracy and the ability to find biologically meaningful interactions for such models. Finally, we evaluated gene characteristics that have an impact on model performance. Gene characteristics such as number of TSSs, gene expression sparsity and exon length lead to differences in performances between single-cell and meta-cell models. Our pipeline is available for the community via the MetaFR repository: https://github.com/SchulzLab/MetaFR.

## Results

### Modeling gene-specific expression with MetaFR

We developed a fully automated Nextflow pipeline, MetaFR, to accurately predict gene expression from single-cell transcriptomic and epigenomic data. An overview of the pipeline is shown in Figure 1. Supplementary Figure 1 provides a detailed flow chart. Each gene is assigned its own model, trained in a gene-specific manner. The features of the gene expression prediction models are defined as equal-sized non-overlapping bins *b*_1_ … *b*_*m*_ that hold the quantified epigenetic signal for each sample *s*_1_ … *s*_*n*_ in a window around a gene (Fig. 1A). Within this study, we use a 1 MB window surrounding the most 5^*′*^ GENCODE ([34]) TSS of a gene, and examine varying bin sizes of 100*bp*, 250*bp*, and 500*bp*. To predict gene expression we chose a Random-Forest (RF) regression approach, given its ability to model non-linear dependencies and to avoid overfitting, while remaining computationally efficient. The models can be trained on single-cells when multiome data is provided. To address the inherent sparsity of single-cell data, the pipeline allows for cells to be grouped into meta-cells based on their gene expression similarities. After generating meta-cells, we assign cells from the scATAC-seq dataset to these RNA meta-cells, resulting in final meta-cells that incorporate both modalities. This enables our method to handle unpaired single-cell data (see Methods), where transcriptomic and epigenomic profiles originate from different cells, provided they have been integrated into a shared feature space beforehand. For model training, the feature matrix is generated for each gene and contains binned activity within the window around the gene, either at the single-cell or meta-cell level. For each gene, the trained RF model for expression prediction and a performance report are provided as output. The report contains the training and test error (Mean Squared Error, MSE) as well as the Pearson and Spearman correlation between predicted and actual expression (Fig. 1B).

**Figure 1.**
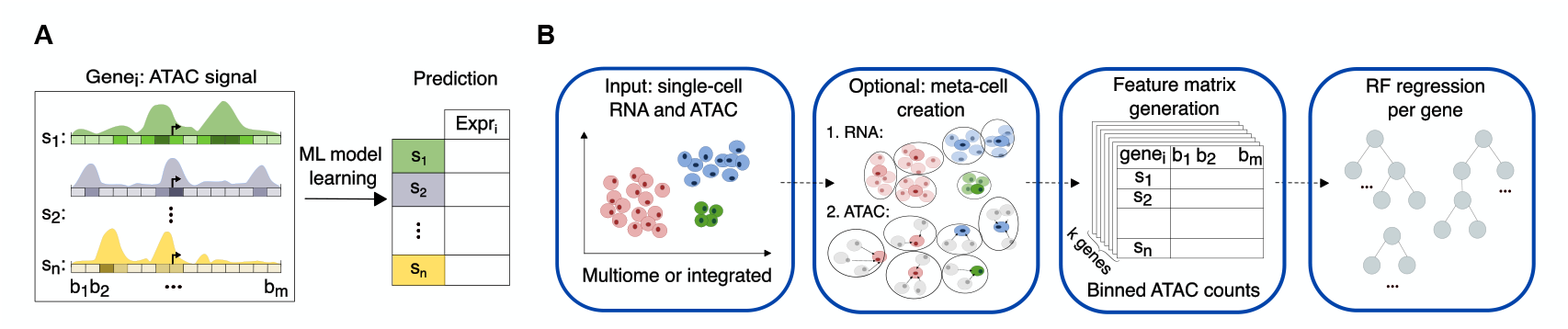
Overview of the MetaFR pipeline. **A** General learning setup: gene-specific models are learned using the epigenetic signal across samples *s*_1_ … *s*_*n*_ in a window around a gene. The epigenetic signal is quantified in equal-sized bins *b*_1_ … *b*_*m*_ and used to predict expression of gene_*i*_ (*Expr*_*i*_). **B** MetaFR is an automated Nextflow pipeline that utilizes scRNA and scATAC data to predict gene expression in a gene-specific manner on single-cell or meta-cell level using Random Forest (RF)-regression trees. For the single-cell setup, multiome data is required. When the meta-cell creation step is applied, unpaired epigenetic and transcriptomic data can be leveraged. In this case, RNA and ATAC cells have to be integrated beforehand to be aligned in a shared feature space.

### Prediction accuracy increases after meta-cell aggregation

We applied our pipeline to a publicly available multiome PBMC dataset from 10x (https://www.10xgenomics.com/datasets/10-k-human-pbm-cs-multiome-v-1-0-chromium-x-1-standard-2-0-0) with approximately 10,000 cells, which is widely used for benchmarking due to its high quality. To assess the model accuracy in an unbiased manner, we randomly held out 20% of the dataset for model testing. The number of input genes for the training phase was 36, 601. We learned for around 19, 300 genes for all tested MetaFR setups. The number of models varies slightly between the methods. Figure 2 shows the prediction performance of different MetaFR setups, evaluated by the Spearman correlation between predicted and observed gene expression on the test set. When investigating the impact of grouping different numbers of cells per meta-cell, we found that increasing the meta-cell size leads to an improved prediction accuracy (Fig. 2A). However, with 150 cells per meta-cell, performance decreases slightly compared to the setting with 100 cells per meta-cell. The decrease can be explained by the number of training instances available after aggregation, as merging more cells into one meta-cell lowers the total number of samples. Since the setup with 100 cells per meta-cell yielded the best performance it was used in all subsequent analyses. A pairwise comparison of the meta-cell setups based on MSE is provided in Supplementary Figure 2A. Figure 2B depicts model performance for genomic feature sizes of 100, 250, and 500 base-pairs for the best performing meta-cell setup. The results for the single-cell setup can be found in Supplementary Figure 2B. Overall, the effect of bin size on performance was minor, although larger bins slightly improved the prediction accuracy. In contrast, the runtime per gene in seconds decreased substantially, from approx. 4.2 for 100*bp* bins to approx. 3 for 500*bp* bins at single-cell level. Consequently, we chose a feature size of 500*bp* for all subsequent analyses.

**Figure 2.**
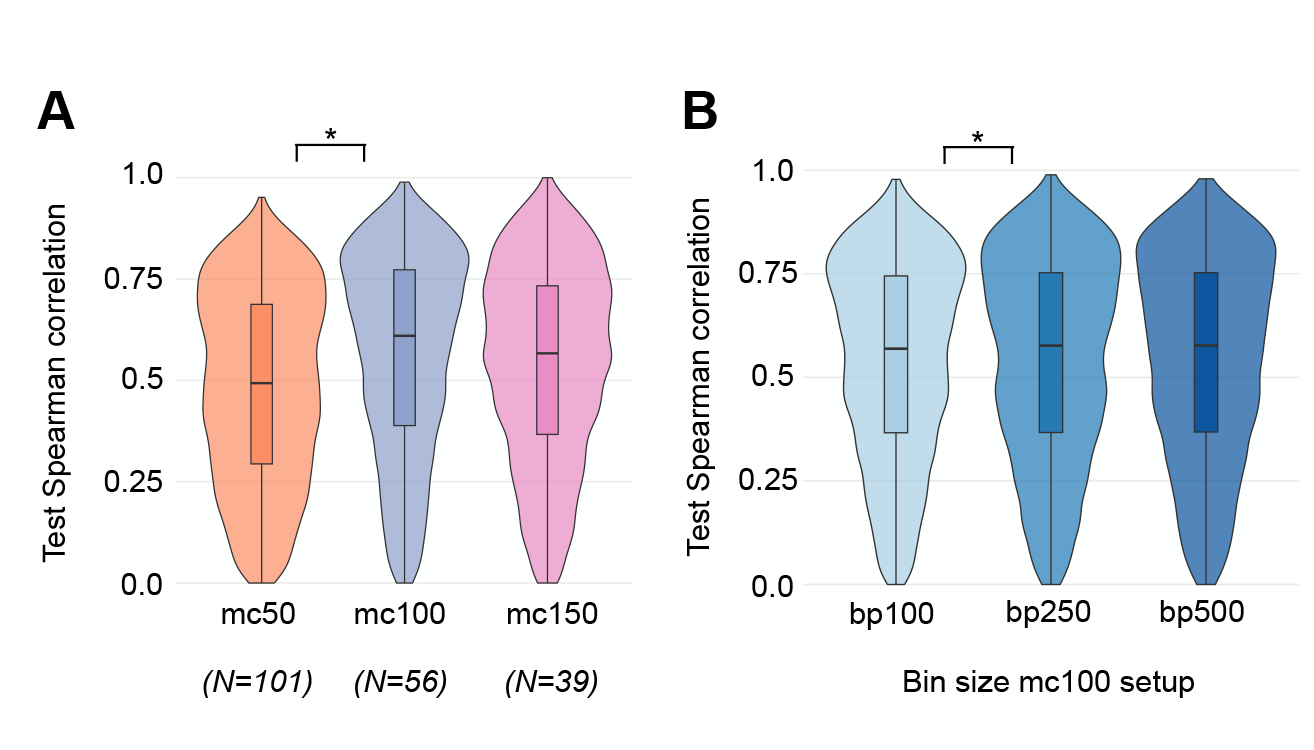
Model performance assessment for meta-cell (mc) MetaFR setups. To assess model accuracy the Spearman correlation was calculated between predicted and actual gene expression on a test set (20% of all data). **A** Correlation on test data for the meta-cell setups with 50, 100 and 150 cells aggregated into one meta-cell for 10, 754 genes, where all setups could obtain a model with test correlation > 0. **B** Correlation on test data for the meta-cell setup with 100 cells per meta-cell and bin sizes varying between 100, 250 and 500 base-pairs. Center line is median, boxlimits correspond to IQR, whiskers to 1.5x IQR. Outliers are not shown. A paired Wilcoxon signed-rank test was performed between gene sets of different setups (p-value ≤ 0.05 indicated with an asterisk).

In order to evaluate the impact of meta-cell generation on the prediction performance, a comparison was made between aggregated single-cell predictions and meta-cell predictions (Fig. 3). To enable direct comparison between meta-cell and single-cell prediction methods, single-cell predictions were summarized into meta-cell representations. More specifically, the single-cell predictions for cells that belong to the same meta-cell were summed. The cell-to-meta-cell assignment was derived from the meta-cell setup. Finally, the Spearman correlation was calculated between predicted and actual gene expression (Fig. 3A). Overall, one can observe that for the majority of genes the meta-cell predictions achieve higher accuracy than the single-cell predictions, although there is a substantial number of genes for which this is not the case (Fig. 3B). For the two setups described above, the number of gene models with a test correlation > 0.5 was 7, 217 for the meta-cell setup and 4, 774 for the single-cell setup (using correlation estimates after aggregation; Fig. 3C). Additionally, the runtime per gene was approximately 0.42 seconds for the meta-cell setup compared to approximately 3 seconds for the single-cell setup (Fig. 3D). We evaluated the differences in the ability between the single-cell and meta-cell setups to identify biologically meaningful region-gene interactions (Fig.4). As ground truth interactions, expression-quantitative trait loci (eQTL) gene interactions were obtained from GTEx [35] for the whole blood tissue. To rank the region-gene interactions obtained from the gene models the feature importance was estimated using absolute SHAPley values ([19]) in a cell type-specific manner (see Methods). We found that, overall, the meta-cell setups achieved significantly higher enrichment than the single-cell setup. Among them, the configuration with 100 cells per meta-cell performed best, capturing the largest number of biologically meaningful interactions compared to the other setups.

**Figure 3.**
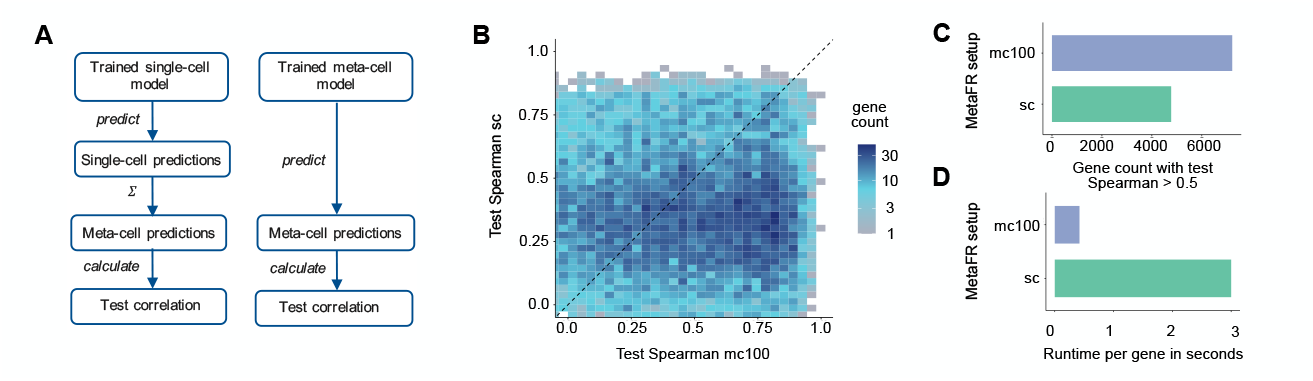
Model performance comparison between single-cell (sc) and meta-cell (mc) MetaFR setups. To assess model accuracy the Spearman correlation was calculated between predicted and actual gene expression on a 20% test set. **A** Workflow for direct comparison of meta-cell and single-cell accuracy: single-cell predictions are aggregated across test cells assigned to the same meta-cell as in the meta-cell setup. **B** Test correlation for meta-cell setup with 100 cells per meta-cell and aggregated single-cell predictions according to A. **C** Number of genes where a model could be generated with test correlation > 0.5. **D** Runtime per gene on an Intel Xeon CPU node with 64 Cores at 2.6-3.3 GHz Frequency and 384 GiB RAM using 25 cores.

**Figure 4.**
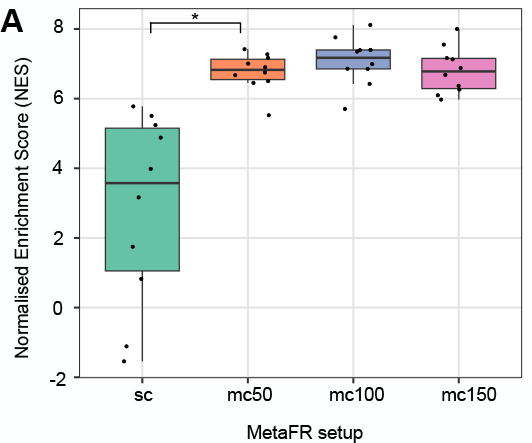
Comparison between single-cell (sc) and meta-cell (mc) MetaFR setups in ability to identify biologically meaningful interactions. **A** Cell type-specific GSEA Normalized Enrichment Score (NES) ([10]) of GTEx ([35]) whole blood eQTLs in the top 100, 000 ranked interactions of the 1, 000 most variable genes. Interactions were ranked across genes using IQR-scaled absolute SHAP values. Additionally, a distance decay weighting was applied ([8]), since eQTLs are biased to be in close proximity to a gene’s promoter ([2, 22]). Cell types with fewer than 100 cells and fewer than 800 eQTL hits among the most variable genes were excluded. The Center line denotes the median, boxlimits correspond to IQR, whiskers to 1.5x IQR. A paired Wilcoxon signed-rank test was performed between gene sets of different setups. P-value ≤ 0.05 indicated with an asterisk).

### Gene characteristics explain performance differences between single-cell and meta-cell models

We evaluated which gene characteristics contribute to the differences in prediction accuracy between single-cell and meta-cell MetaFR (100 cells per meta-cell) (Fig. 5). Genes included in the analysis were those with aggregated single-cell (see Methods) or meta-cell test Spearman correlations ≥ 0.3, and where one setup outperformed the other by at least 0.1 correlation. Figure 5A indicates that single-cell MetaFR performs better for highly sparse genes. Analysis of the number of annotated GENCODE ([34]) TSSs shows that the single-cell setup outperforms meta-cell MetaFR for genes with fewer TSSs and shorter exon lengths. Furthermore, single-cell models tended to perform better for genes with fewer annotated GENCODE [34] transcripts, shorter gene length, and lower number of genes within a 1 MB window (Suppl. Fig. 3A). Genes were stratified into performance groups according to the test correlation, revealing consistent trends across meta-cell and single-cell setups, except that longer genes and denser loci improved meta-cell performance but reduced single-cell performance (Suppl. Fig. 3B+C).

**Figure 5.**
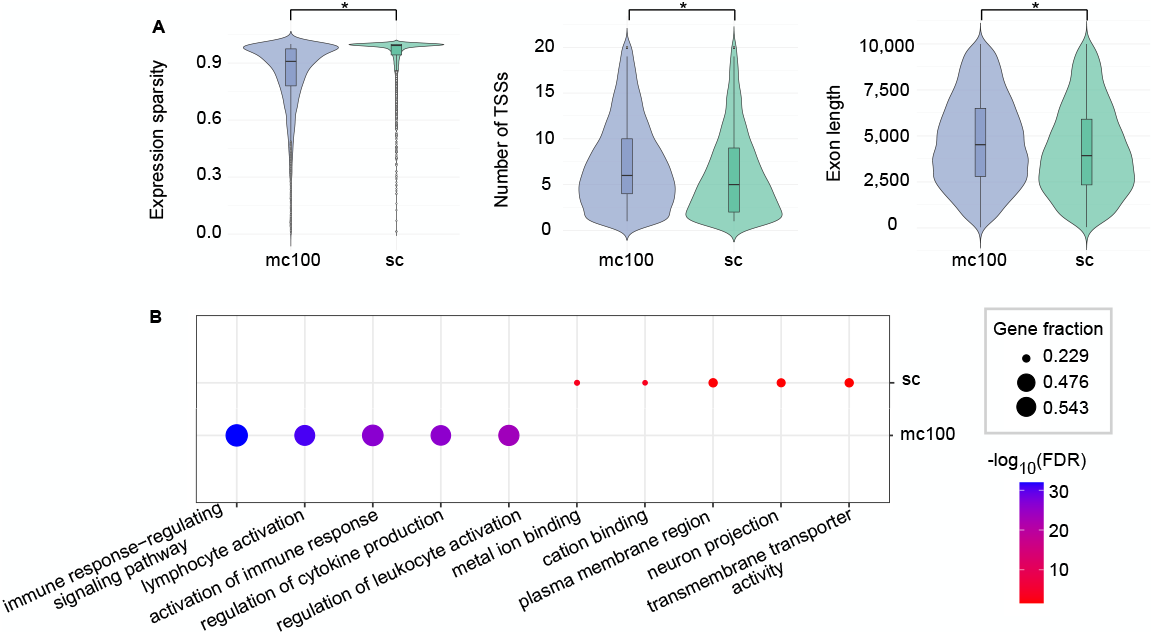
Investigation of gene characteristics that impact performance differences between single-cell (sc) and meta-cell (mc) setups. The best-performing gene sets for the single-cell (4, 615 genes) and meta-cell setup with 100 cells per meta-cell (7, 172 genes) are defined as the models that achieve a minimum performance of 0.3 test correlation and where one method outperforms the other by > 0.1. **A** Assessed gene characteristics include gene expression sparsity, corresponding to fraction of 0 in the expression in test cells, number of annotated GENCODE ([34]) TSSs and total exon length. (Center line is median, box limits correspond to IQR, whiskers to 1.5x IQR. An unpaired Mann-Whitney U Test was performed between the best-performing gene sets. P-value ≤ 0.05 indicated with an asterisk). **B** GO enrichment with g:Profiler ([18]) of best-performing gene-sets. The background was set to the initial gene set of 36, 601 genes. High-level terms (depth ≤ 7) were filtered with R Package ontologyIndex ([12]). Displayed GO terms were selected from the top 100 significant terms. All terms can be found in Supplementary Material.

A GO enrichment analysis of the genes that performed best in the single-cell setup identified a total of 15 significant terms. A representative selection of these terms is shown in Fig. 5B. The enriched terms predominantly relate to basic cellular processes, reflecting the core functions captured by the single-cell models. In contrast, 8, 804 GO terms were enriched among the best-performing meta-cell genes. Notably, within the top 100 most significant biological process terms, several were cell-type-specific, including processes such as activation of the immune response.

### MetaFR outperforms the state-of-the art method SCARlink in accuracy and runtime

We compared MetaFR to the state-of-the-art multi-variate regression approach SCARlink ([21]) regarding the prediction accuracy at single-cell level. The feature setup is identical for both approaches: The epigenetic signal is binned with 500 bp in a window around the target gene. For SCARlink the size of the genomic window varies between genes, since a 250 kb extension upstream and downstream of the gene body is applied. It uses *L*_1_-regularized Poisson regression as a prediction method. All subsequent analyses are conducted at the single-cell level. To assess the prediction accuracy we calculated the MSE and Spearman correlation between predicted and actual gene expression on the same test cells. Following the default practice for SCARlink, the input gene set was defined as the 2, 000 most variable genes. A model could be generated for 536 genes. The number of genes where both approaches obtained a model was 495. Regarding the MSE, MetaFR outperforms SCARlink on 354 of those genes (Fig. 6A). The number of genes where a method was able to generate a model with test correlation above 0.3 amounts to 631 for MetaFR and 316 for SCARlink, respectively (Fig. 6B). SCARlink required approximately 4.3 minutes to learn one gene, whereas MetaFR required approximately 3 seconds.

**Figure 6.**
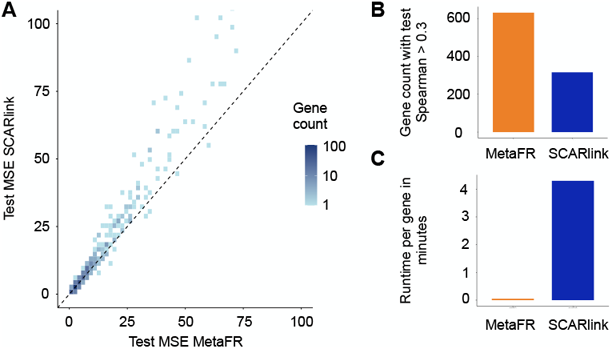
Comparison to state-of-the-art method SCARlink. **A** Test error (MSE) of MetaFR and SCARlink for 495 genes, for which both methods produced a model. **B** Number of genes where a model could be generated with test correlation (Spearman) > 0.3 from their respective initial gene sets (see Methods). **C** Runtime per gene on an Intel Xeon CPU node with 64 Cores at 2.6-3.3 GHz Frequency and 384 GiB RAM using 25 cores. The test error and correlation are calculated between predicted and actual gene expression.

## Discussion

We developed MetaFR, a fully automated Nextflow pipeline that generates gene-specific regression-based Random Forest models to predict gene expression from single-cell RNA-seq and ATAC-seq data. Our pipeline allows time-efficient analysis and obtains reliable models of gene expression, which can be used to study gene regulatory elements in any organism for which single-cell RNA-seq and ATAC-seq data is available. We were able to outperform the state-of-art gene-specific multivariate method for gene expression prediction SCARlink ([21]) in both prediction accuracy and runtime. We systematically evaluated the impact of meta-cell creation on expression prediction accuracy. Our analysis revealed that meta-cell aggregation leads to a significant increase in model accuracy. Although increasing the number of cells per meta-cell generally improves the performance, it is essential to retain a sufficient number of training samples. Overall, the choice of meta-cell size is highly dataset specific. It depends on factors such as number of cells per cell type or experimental conditions. A particular challenge is the presence of rare cell types, which are often aggregated into a single meta-cell, limiting their contribution to model training. In addition, meta-cell generation is strongly influenced by the quality of the integration. To mitigate this impact, metadata such as cell type can be incorporated in the clustering process.

An advantage of using Random Forest models for prediction of gene expression is their computational efficiency, which facilitates the exploration of different parameter settings, particularly tuning the number of cells per meta-cell. Although overall performance improved with meta-cell aggregation, a substantial subset of genes was better predicted by single-cell models. These genes were lowly expressed, with a median sparsity of around 95%. A possible explanation is that, for highly sparse genes, meta-cell summarization overly reduces expression variability, thereby limiting the model’s ability to learn informative patterns.

We found that single-cell models perform better for genes with simpler structure (shorter exon lengths and fewer TSSs). Correlation analysis indicated a negative association between the expression sparsity and exon lengths (≈ − 0.39), as well as expression sparsity and number of TSS (≈ − 0.44). Since there are fewer regions to sample during sequencing of shorter exons, sparsity naturally increases.

In addition, RF models have difficulty predicting exact zeros because of the averaging across trees. In the future, as larger and higher-resolution datasets become available, models can be trained at the single-cell level to fully leverage expression variability between individual cells. More sophisticated machine learning methods, such as neural networks, which currently face challenges for gene-specific single-cell models due to their large number of hyperparameters, could be applied. However, these models are considerably more computationally intensive, which is an important consideration given that tens of thousands of models need to be trained. With MetaFR, we provide a highly efficient pipeline for accurate gene expression prediction, including an optional aggregation of individual cells into meta-cells to enhance signal, making it well-suited for application to large-scale datasets in future studies.

## Methods

### Data pre-processing

We used the multiome 10x Genomics PBMC dataset^1^. We downloaded the filtered feature barcode matrix, the ATAC per fragment information, and index files. We applied the filtering steps from the Seurat weighted-nearest-neighbor analysis vignette (https://satijalab.org/seurat/articles/weighted nearest neighbor analysis): all cells with scATAC counts> 70, 000 and < 5, 000, and with scRNA counts < 1, 000 and > 25, 000 were filtered. In addition, cells with more than 20% mitochondrial reads were excluded, resulting in 10, 412 high quality cells. The gene expression was CPM normalized (scaling factor 10, 000). ATAC and RNA cell integration, was done using the Seurat vignette for that purpose (https://satijalab.org/seurat/articles/seurat5 atacseq integration vignette). The main steps included computing gene activity scores (GeneActivity() function), using them to identify anchors (mutual nearest neighbors via FindTransferAnchors()) for CCA-based integration, and finally co-embedding RNA and ATAC cells with TransferData(). Seurat ([14]) v4.3.0 was used.

### MetaFR pipeline

The subsequent steps have been fully automated with Nextflow (version 25.04.6). The required input files for our pipeline are a fragment file containing the fragment positions, barcodes and the total read count and the index file for the epigenetic modality, and the gene expression count matrix. In addition, a Seurat object containing the integrated RNA and ATAC cells is needed when applying the meta-cell aggregation step.

#### Meta-cell creation

For generating the optional meta-cells, a Seurat object containing the integrated RNA and ATAC cells is required as input. Here, single-cell RNA-seq and ATAC-seq data can be either measured in the same cell (multiome) or different cells (unpaired). Two main steps were performed for signal summarization: first, the cells containing the RNA-seq signal were clustered using a k-medoid approach based on the similarity of their gene activity profiles. The cluster centers represent the RNA meta-cells. Next, the cells containing the epigenetic signal were assigned to the closest RNA meta-cell with a k-nearest-neighbor (kNN) approach. For both steps, the Euclidean distance was applied as the distance metric. For our analyses, the parameter *k*, corresponding to the number of cells per meta-cell, was varied between 50, 100 and 150. Varying this parameter impacts the total number of meta-cells. Our implementation is available as an R package **MetaCellaR**: https://github.com/fba67/MetaCellaR.

#### Meta-cell filtering

For all analyses, we filtered meta-cells with less than 200, 000 ATAC-seq counts.

#### Activity Binning

Prior to model learning, the feature matrices were generated for each gene. They contain the ATAC signal quantified within equal-sized bins *b*_1_ … *b*_*m*_ in a 1 MB window centered on the most 5^*′*^ GENCODE ([34]) TSS of a gene for each sample *s*_1_ … *s*_*n*_ (single-cell or meta-cell). We varied the bin sizes between 100, 250 and 500 basepairs. The ATAC activity was TF-IDF normalized and for activity binning, the Signac ([32]) FeatureMatrix function was used.

#### Random Forest Regression

For the model training we applied a Random Forest regression approach on the binned epigenetic data with the `RandomForestRegressor` function from the scikit-learn library in Python. The response was min-max scaled to the range [0,1]. The default parameter settings were used for all analyses:

~~~
n_estimators=100, criterion=‘squared_error’,max_depth=None,
min_samples_leaf=1, min_weight_fraction_leaf=0.0,
max_features=1.0, max_leaf_nodes=None,min_samples_split=2,
min_impurity_decrease=0.0, bootstrap=True,monotonic_cst=None,
oob_score=False, n_jobs=None,random_state=0, verbose=0,
warm_start=False, ccp_alpha=0.0,max_samples=None
~~~

### Performance assessment

For the single-cell and meta-cell setups we randomly held-out 20% of the cells or meta-cells for model testing. All 19 cell types are present in both, train and test set for the single-cell setup. For the meta-cell setups, 18 cell types are present in the training set, CD4 Naive meta-cells are only present in the test set. To evaluate prediction accuracy, the Spearman correlation, as well as the Mean-Squared-Error (MSE) was calculated between predicted and actual gene expression for each setup on the corresponding test set.

### eQTL enrichment analysis

To evaluate differences in meta-cell and single-cell setups in their ability to capture interactions supported by eQTL gene-pairs on the PBMC dataset, the *Whole Blood* tissue from GTEx was chosen. The fine-mapping method with the highest number of eQTL-gene pairs, DAP-G, was selected (‘GTEx v8 finemapping DAPG.CS95.txt.gz’ file). The 1, 000 genes with the highest variability in the observed expression were considered for this analysis. For each MetaFR setup (50, 100 and 150 cells per meta-cell and single-cell) we calculated approximated SHAPley values using the TreeSHAP ([19]) algorithm (shap library version 0.47.1) to obtain the regions that are important for a model’s prediction. To compare single-cell and meta-cell setups, the mean SHAP value for each cell type was calculated. To rank the genomic bin-gene interactions across genes, the absolute SHAP values were scaled to interquantile range (IQR) using the RobustScaler function from the scikit-learn library in Python. In addition, the IQR-scaled SHAP values *shap*_*g,b*_ for each interaction *g, b* between gene *g* and bin *b* were weighted with a distance-decay function ([8]):

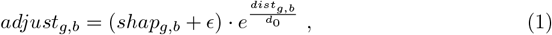

where *ϵ* was set to 0.05, *dist*_*g,b*_ represents the distance between a gene’s most 5^*′*^ TSS and genomic bin, and weight decay scaling parameter *d*_0_ = 200, 000bp. Then, the normalized enrichment score of eQTLs in the top 100, 000 highest ranked interactions was calculated using the GSEApy ([10]) pre-rank function (v.1.1.2) in a cell type-specific manner. The number of permutations was set to 100. The weighting parameter p was set to zero. In order to ensure the same number of interactions supported by eQTL-gene pairs for each cell type, the number of overlapping eQTL-gene pairs was reduced to the minimum overlap across all cell types.

### Performance comparison MetaFR and SCARlink

SCARlink ([21]) employs a 250kb window around the gene body for each gene. The size of the window surrounding the most 5^*′*^ GENCODE TSS of a gene was set to 1 MB for MetaFR. The epigenetic signal was quantified in 500 bp bins for both methods. The gene expression was CPM normalized with scaling factor 10, 000. Following the default practice for SCARlink, the input gene set was defined as the 2, 000 most variable genes. For MetaFR, all 36, 601 genes present in the dataset without any filtering on the expression variance were used for model training. For SCARlink, the ArchR ([11]) AddTileMatrix function was used for generating the feature matrices. All analyses were performed on single-cell level on the same set of test cells. 20% of the data set was randomly selected for testing.

## Supporting information

GO terms

## Data availability

We downloaded the PBMC dataset from the 10x Genomics Website: https://www.10xgenomics.com/datasets/10-k-human-pbm-cs-multiome-v-1-0-chromium-x-1-standard-2-0-0. The pipeline and manuscript code are publicly available on GitHub: https://github.com/SchulzLab/MetaFR.

## Author contributions statement

L.R. implemented the MetaFR pipeline, designed and conducted the experiments, wrote the paper. F.B.A. designed and implemented the MetaCellar R package, D.H. helped with analysis, M.H.S. supervised the work, designed experiments, acquired funding. All authors proofread the final manuscript.

## Acknowledgments

This work was supported by the Goethe University Frankfurt am Main, the German Centre for Cardiovascular Research (DZHK Standort Rhine Main 81Z0200101 to M.H.S., L.R. and D.H.), the Deutsche Forschungsgemeinschaft (DFG) excellence cluster EXS2026 (Cardio-Pulmonary Institute, project-ID 390649896 to M.H.S.), the DFG project-ID 403584255 - TRR 267 (TP Z03 to D.H., F.B.A. and M.H.S.), and DFG project-ID 456687919 - SFB1531 (TP S03 to L.R. and M.H.S.). M.H.S. acknowledges the Hessian.AI for funding.

## Footnotes

1 https://www.10xgenomics.com/datasets/10-k-human-pbm-cs-multiome-v-1-0-chromium-x-1-standard-2-0-0

