## Supplementary Materials for "Predicting gene-specific regulation with transcriptomic and epigenetic single-cell data"

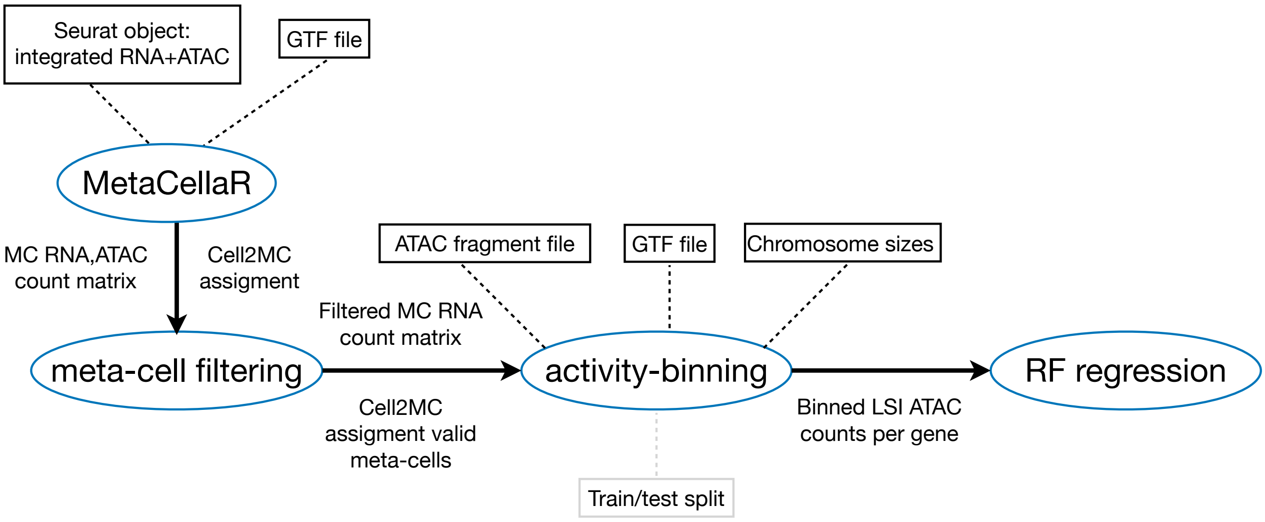


**Fig. 1. Flowchart MetaFR Nextflow pipeline.** MetaCell creation (MetaCellaR and meta-cell filtering) is optional. For MetaCellaR a Seurat object containing the integrated RNA and ATAC cells is required as input together with a GTF file. The output is the gene expression and ATAC count matrix on meta-cell level, as well as the assignment of cells to meta-cells. In the meta-cell filtering step, meta-cells that contain less than $200,000$ ATAC reads are excluded. This threshold can be adjusted by the user. For the activity-binning step, additional input files are needed: the ATAC fragment file and chromosome sizes. Optionally, a train-test partition of the cells/meta-cells can be provided. If not, a random train-test split (default = 20%) will be applied. The LSI-normalized ATAC count matrices are then used to train a RF regression model per gene.

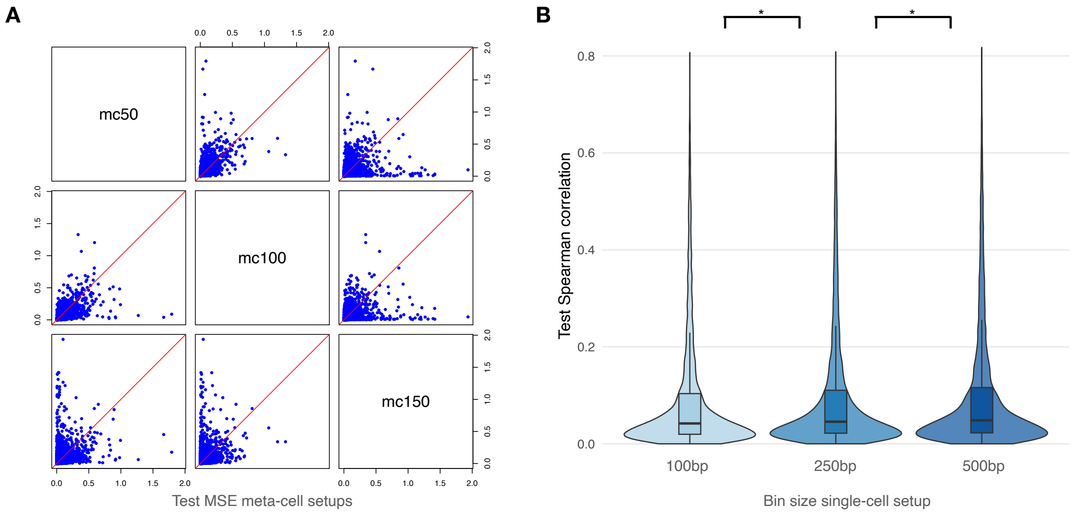


**Fig. 2. Model performance assessment for meta-cell (mc) and single-cell (sc) MetaFR setups.** To assess model accuracy the Spearman correlation was calculated between predicted and actual gene expression on a test set (20% of all data). **A** MSE on test data for the meta-cell setups with 50, 100 and 150 cells aggregated into one meta-cell for $19,306$ genes. The MSE was calculated on the log-transformed CPM expression values. **B** Correlation on test data for the single-cell setup and bin sizes of 100, 250 and 500 base-pairs. Center line is median, boxlimits correspond to IQR, whiskers to 1.5x IQR. Outliers are not shown. A paired Wilcoxon signed-rank test was performed between gene sets of different setups (p-value $\leq 0.05$ indicated with an asterisk).

**Fig. 3. Investigation of gene characteristics that impact model performance.** Assessed gene characteristics are gene length, gene density ($=$ number of genes within the $1$ MB window around the gene), number of annotated GENCODE transcripts, gene expression sparsity (fraction of zeros in expression across test cells), number of annotated GENCODE TSSs, and total exon length. **A** The best-performing gene sets for the aggregated single-cell sc (4,615 genes) and meta-cell mc100 setup with 100 cells per meta-cell (7,172 genes) are defined as the models that achieve a minimum performance of 0.3 test correlation and where one method outperforms the other by $> 0.1$. **B+C** Gene sets were classified based on their test correlation for single-cell (not aggregated) and meta-cell ($100$ cells per mc): $< 0.1$ is defined as failed, $< 0.5$ as medium and $> 0.5$ as high. **B** MetaFR setup with 100 cells per meta-cell with $7,106$ high performing, $5,536$ medium performing and $6,111$ failed genes. **c** Single-cell MetaFR with $159$ high performing, $1970$ medium performing and $16,319$ failed genes. Center line is median, box limits correspond to IQR, whiskers to $1.5\times$ IQR. An unpaired Mann–Whitney U test was performed between the best-performing gene sets. P-value $\leq 0.05$ indicated with an asterisk.

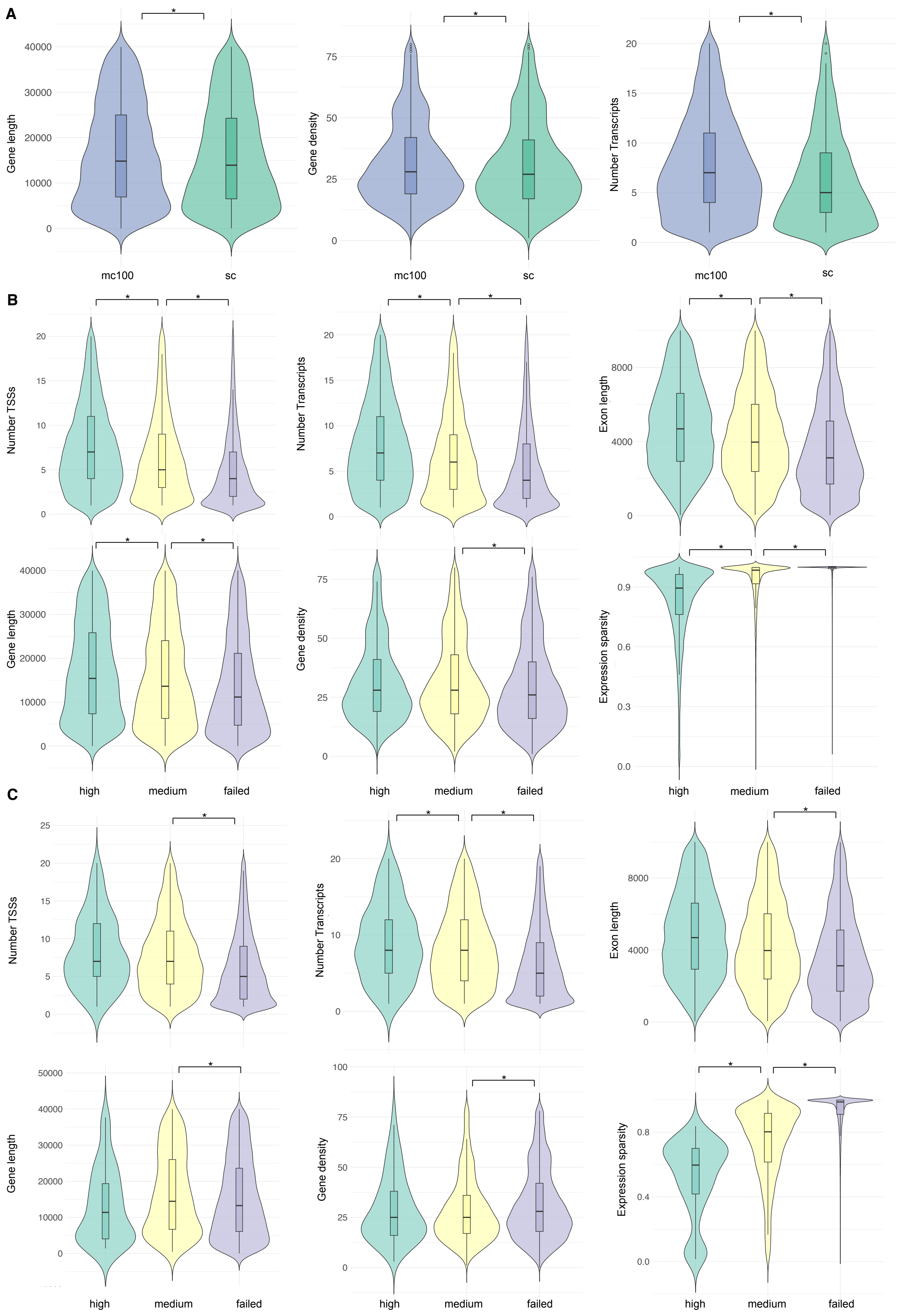
A
Gene length
40000
30000
20000
10000
0
mc100
sc
Gene density
75
50
25
0
Number Transcripts
20
15
10
5
0
B
Number TSSs
Number Transcripts
Exon length
8000
4000
Gene length
Gene density
Expression sparsity
0.9
0.6
0.3
0.0
high
medium
failed
C
Number TSSs
Number Transcripts
Exon length
Gene length
50000
Gene density
100
Expression sparsity
0.8
0.4
0.0
*

### Supplementary Files

The following supplementary files are provided:

- **All GO terms for MetaFR with 100 cells per meta-cell** `go_enrichment_mc100_spearman_q0_500bp_all_terms.tsv` — full list of 8,804 significant Gene Ontology terms.
- **All GO terms for single-cell MetaFR** `go_enrichment_aggr_sc_spearman_q0_500bp_all_terms.tsv` — full list of 15 significant Gene Ontology terms.
